# Performance Evaluation of Empirical Mode Decomposition Algorithms for Mental Task Classification

**DOI:** 10.1101/076646

**Authors:** Akshansh Gupta, Dhirendra Kumar, Anirban Chakraborti, Kiran Sharma

## Abstract

Brain Computer Interface (BCI), a direct pathway between the human brain and computer, is one of the most pragmatic applications of EEG signal. The electroencephalograph (EEG) signal is one of the monitoring techniques to observe brain functionality. Mental Task Classification (MTC) based on EEG signals is a demanding BCI. Success of BCI system depends on the efficient analysis of these signals. Empirical Mode Decomposition (EMD) is a filter based heuristic technique which is utilized to analyze EEG signal in recent past. There are several variants of EMD algorithms which have their own merits and demerits. In this paper, we have explored three variants of EMD algorithms named Empirical Mode Decomposition (EMD), Ensemble Empirical Mode Decomposition (EEMD) and Complete Ensemble Empirical Mode Decomposition with Adaptive Noise (CEEMDAN) on EEG data for MTC-based BCI. Features are extracted from EEG signal in two phases; in the first phase, the signal is decomposed into different oscillatory functions with the help of different EMD algorithms and eight different parameters (features) are calculated for each function for compact representation in the second phase. These features are fed into Support Vector Machine (SVM) classifier to classify the different mental tasks. We have formulated two different types of MTC, the first one is binary and second one is multi-MTC. The proposed work outperforms the existing work for both binary and multi mental tasks classification.

## 1 Introduction

Human brain has the capability to distinguish two or more different tasks without much effort. In literature, most of the research works have been suggested to distinguish between two different tasks at a given instant of time; a few research works deal with multi mental task classification (Anderson et al., 2011; Donoghue, 2002; Li et al., 2014; Palaniappan et al., 2002; Wang et al., 2012; Zhang et al., 2010). There is a need of a multi mental task classification system that can distinguish more than two mental tasks at a given instance of time. As the number of chosen classes grows, it becomes more difficult to classify a test sample correctly. The computational complexity of the multi class problem is much higher in comparison to a binary class problem.

The electroencephalograph (EEG) signal is one of the monitoring techniques to capture brain activity corresponding to a given task based stimulation. The amplitude of the captured EEG signals is low. Hence, the signal in its raw form is not helpful to distinguish among different mental tasks at a given time. Given these facts, classification of multi mental tasks is considered to be a challenging problem. However, limited BCI models (Li et al., 2014; Palaniappan et al., 2002; Zhang et al., 2010) have been proposed to distinguish more than two tasks at a given instance of time. Therefore in this study, we have formulated a problem for the multi mental task as well as binary mental task classification with the help of EEG signals. One versus rest approach based support vector machine (SVM) is used as a multi mental task classifier to build the decision model. The overall flow chart of proposed model has been shown in Figure 1. Rest of the paper is organized as follows: In section 2, the state of art of feature extraction for BCI as well as multi-class BCI is given. Section 3 contains the description of proposed approach. Experimental data and the related discussion are given in section 4, and finally section 5 draws the conclusion.

**Figure 1:**
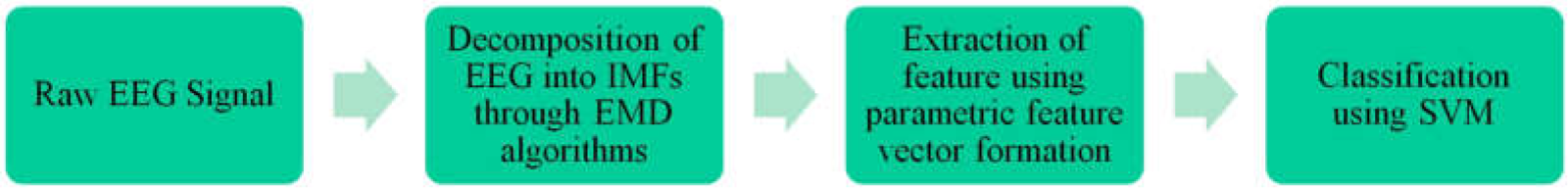
Schematic flow chart of the propo ed model for Men al Task Classification

## 2 Related Works

In literature, various feature extraction techniques have been studied and suggested for BCI (Bashashati et al., 2007). These feature extraction techniques can be grouped into three major categories: (i) Temporal methods (ii) Frequency domain methods and (iii) hybrid of temporal and frequency domain methods.

The temporal methods are predominantly adaptive to describe neurophysiological signals with an accurate and specific time information. The temporal variations of the signal are characterized by the extracted features. In time domain, amplitude of the signal or statistical measures like absolute mean, standard deviation and kurtosis of the signal are used to characterize EEG signal(Bostanov, 2004; Hjorth, 1970; Motamedi-Fakhr et al., 2014; Vidaurre et al., 2009).

It is known that EEG signals consist of a set of explicit oscillations, which are known as rhythms. Corresponding to different mental tasks, different rhythms are associated with these EEG signals (Canolty and Knight, 2010; Keren et al., 2010; Klimesch, 2012; Pfurtscheller and Da Silva, 1999; Sauseng et al., 2010). Hence, frequency information can be used as a feature of the signal. These frequency information embedded in the signal is utilized to represent the signal more accurately. Power spectral analysis (density) has been used in literature to extract accurate frequency content features and produce high frequency resolution (Palaniappan et al., 2002).

However, the neurophysiological signal used in BCI have generally specific properties in both the temporal and frequency domain. Frequency spectrum of the EEG signal is observed to vary over time, indicating that the EEG signal is non-stationary in nature. Short-time Fourier transform and wavelet transform are suggested methods to extract both frequency and temporal information based features from nonstationary signal. Such methods for representation of the signal can capture sudden temporal variations in the EEG signal. The Wavelet Transform (WT) (Da ubechies, 1990; Mallat, 1989) is an effective technique which allows analysis of both time and frequency contents of the signal simultaneously. WT is utilized in analysis of EEG signals in the fields of motor imagery and epileptic seizures, (Bostanov, 2004; Cvetkovic et al., 2008; Hsu and Sun, 2009; Ocak, 2009), brain disorders, (Hazarika et al., 1997), classification of human emotions (Murugappan et al., 2010), and non-motor imagery (Cabrera et al., 2010). However, **WT** uses some fixed basis functions which makes it non-adaptive (Huang et al., 1998) to the signal to be processed. Another method for analyzing signals like EEG is Empirical Mode Decomposition (EMD)(Huang et al., 1998),which is a data driven approach. This method is self-adaptive according to the signal to be processed unlike to WT, where a fixed set of basis functions is used. It decomposes a signal into finite, well defined, low frequency and high frequency components known as Intrinsic Mode Functions (IMFs) or modes. The EMD method has been used to extract representative data for BCI (Diez, Mut, Laciar, Torres and Avila, 2009; Gupta et al., 2015; Kaleem et al., 2010) to classify mental task.

This work explores the suitability of EMD and its variants to analyze the EEG for binary as well as multi mental tasks classification problem. A non-parametric statistical test is also carried out to validate the experimental findings.

## 3 Proposed Approach

In this work, features are extracted from EEG signal in two steps: In the first phase, EEG signal corresponding to a given channel is decomposed by Empirical Mode Decomposition (EMD) algorithms. In the second phase, statistical and uncertainty parameters are calculated from each decomposed signal for a given channel, to represent the signal more compactly. Brief description of the variants of EMD are discussed below.

### 3.1 Empirical Mode Decomposition (EMD)

EMD is a mathematical tool which is utilized to analy ze a non-stationary and nonlinear signal. Under the assumption that any signal contains a series of different intrinsic oscillation modes. EMD is used to decompose an incoming signal into its different Intrinsic Mode Functions (IMF). An IMF is a continuous function which satisfies the following conditions (Huang et al., 1998):

1. The number of extrema and the number of zero crossings are either equal, or differ at most by one.
2. The mean value of the envelope defined by the local maxima and local minima is zero at a given point.

The first condition implies that there is need of a narrow band requirement for a signal to be a stationary Guassian process. The second condition is needed for abstaining instantaneous frequency from unwanted fluctuations induced by asymmetric waveforms. The basic steps of EMD are given in Algorithm 1. Figure 2 showed the plot of first four IMFs of a EEG segment using EMD algorithm. From Figure 2 it can be noted that these IMFs can characterized the signal well.

**Figure 2:**
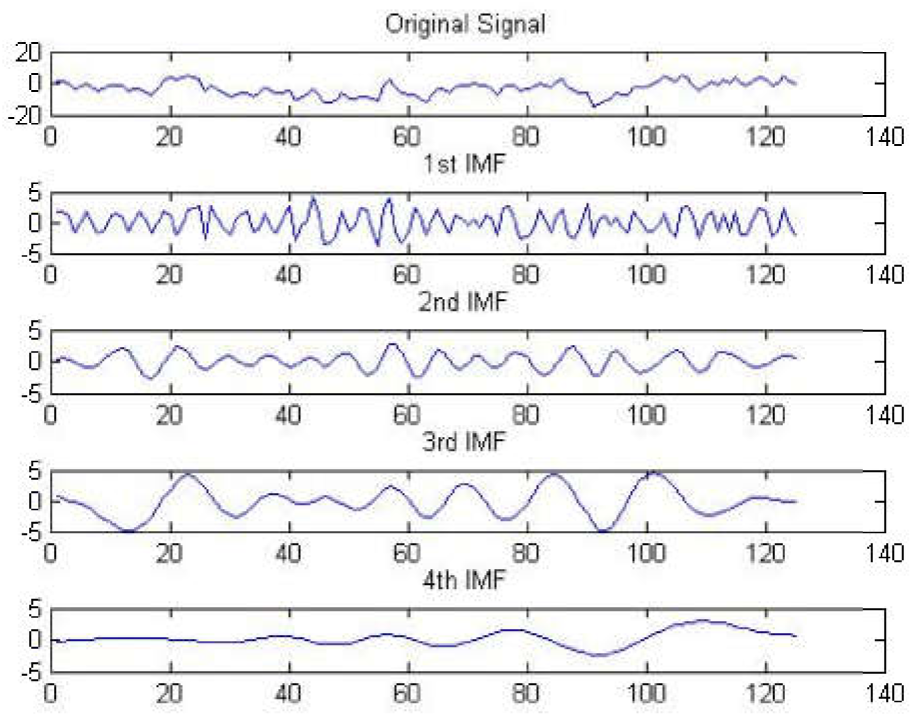
IMF plot obtained for a given EEG signal.

#### Algorithm 1: Algorithm for EMD

1. **Input:** Signal *x(m);*
2. For the signal, *x(m),* identify all local maxima and minima;
3. Calculate the upper envelope by connecting all the local maxima points of the signal using a cubic spline;
4. Repeat the same for the local minima points of the signal to find the lower envelope;
5. Calculate the mean value of both envelopes, say *m*_1_;
6. Update the signal, *x(m)* = *x(m)* − *m*_1_;
7. Continue the steps 2 to 6, and consider *x(m*) as the input signal, until it can be considered as an *IMF* as per the definition stated above;
8. The residue *r*_1_ is obtained by subtracting the first *IMF (IMF*_1_) from *x(m)* i.e. *r* _1_ = *x*(*m*) – *IMF*_1_. The residual of this step becomes the signal *x*(*m*) for the next iteration;
9. Iterate steps 2 to 8 on the residual *r*_*j*_; *j* = l, 2, 3, …, *k* in order to find all the IMFs of the signal until residue *r*_*k*_, becomes a monotonic function from which no more IMF can be extracted.

Thus, a signal *x(m),* can be represented as:

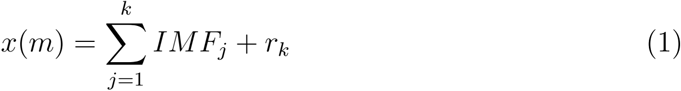

According to Huang et al. (1998), there is one stopping criteria in *T* steps to further produce IMFs based on standard deviation, can be defined as

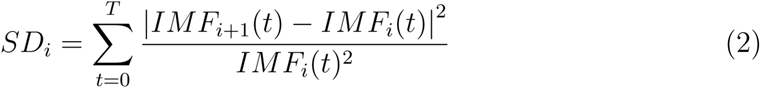

The decomposition process stops when the value of *SDs* is smaller than predefined value.

### 3.2 Ensemble Empirical Mode Decomposition (EEMD)

One of the major problem with EMD method is frequent mode mixing. This problem arises when a single IMF contains signal of widely different scale or a signal of same scale obtained from different IMFs. To alleviate the problem of scale separation, Wu and Huang (2009) have proposed a noise-assisted data analysis method, called Ensemble Empirical Mode Decomposition (EEMD). EEMD defines true IMF components as the mean of an ensemble of the trails which consists of signal plus white noise with finite amplitude (Wu and Huang, 2009). Thus the signal *x(m)* in *i*^*th*^ trial after adding white noise can be represented as

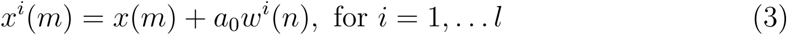

where *w*^*i*^ *(n)* is the white noise in *i*^*th*^ trial with unit variance and *a*_*0*_ amplitude. For each *i* = 1, 2 …*l* the 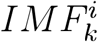 is calculated with different realization of white noise with the signal obtained using Equation 3. The average k^*th*^ 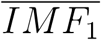 can be defined as

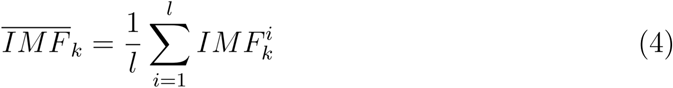

The pragmatic concepts of EEMD are as follows:

1. The added collection of white noise cancels each other with the help of ensemble mean, thus only signal can be one ingredient of the mixture of the signal and white noise.
2. To search all possible solution, it is necessary to ensemble white noise of finite amplitude with signal.
3. To obtain true and physically meaningful IMFs from EMD, it is necessary to add noise to the signal.

### 3.3 Complete Ensemble Empirical Mode Decomposition with Adaptive Noise (CEEMDAN)

The problem of mode mixing in original EMD algorithm is successfully addressed by EEMD by adding white noise into the signal, but this also leads to a problem that noise is not fully segregated from the signal and the resultant different IMFs may contain mixture of noise and signal. To resolve this problem, Torres et al. (2011), have proposed CEEMDAN algorithm that provides good spectral separation of the modes. Hence, It gives an exact reconstruction of the original signal with a lower computational cost.

The first residue can be calculated as:

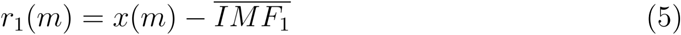

where 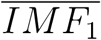 is the first average *IMF* obtained by EEMD. The second average *IMF* can be found as:

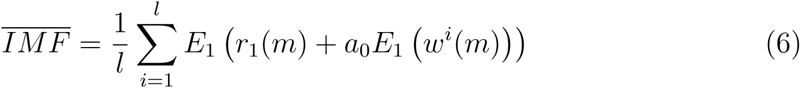

After finding *k*^*th*^ residue, for *k=* 2, …, *K,* the *k*+ 1 average *IMF* can be defined as:

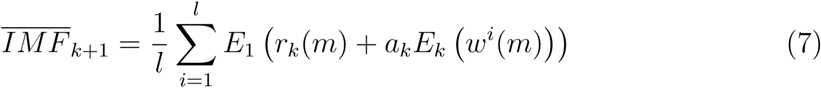

where *E*_*k*_ (.) is an operator to extract *k*^*th*^ IMF from given signal by EMD algorithm and amplitude *a*_*k*_ allow to select the SNR at each stage. Detailed description can be found in Torres et al. (2011).

## 4 Experimental Setup and Result

### 4.1 Dataset and constructing feature vector

In order to compare the efficacy of these EMD algorithms for mental task classification experiments were performed on a publicly available EEG dataset. We have also Compared the proposed model with the work of (Zhang et al., 2010) on the same dataset for multi-mental task classification. This dataset consists the recordings of EEG signals using seven electrode channels (namely C3, C4, P3, P4, 01, 02 and EOG) from seven subjects with the recording protocols described below. Each subject was asked to perform five different mental tasks as: *Baseline task* (relax: *B);* mental *Letter Composing task (L);* Non trivial *Mathematical task (M); Visualizing Counting* (C) of numbers written on a blackboard and *Geometric Figure Rotation (R)* task. Each of the recording session consists of five trials of each of the five mental tasks.

Each trial is of 10sec duration recorded with a sampling frequency of 250 Hz, which resulted into 2500 samples points per trial. We have utilized data of all subjects except Subject 4, due to some missing and incomplete information Faradji et al. (2009). Detailed explanation can be found in the work of Keirn and Aunon (1990)^1^. Six electrodes placed on the scalp at C3, C4, P3, P4, 01 and 02 are used for extracting the feature for mental task classification as EOG gives only artifact. For feature construction, the data of each task of each subject is sampled into half-second segments, yielding 20 segments (signal) per trial for each subject as some researchers have done (Palaniappan et al., 2002). The complete pipeline for constructing the feature vector from each subject using all trial corresponding to each mental task labels (B, L, M, C and R) is described below:

1. The EEG signal corresponding to each task of a given subject is sampled into half-second segments, yielding 20 segments (signal) per trial per channel.
2. In this way, corresponding to each channel, each of the 20 segments are used to generate the 4 IMFs using EMD algorithms.
3. To represent each of these IMFs per segment per channel compactly, eight statistical or uncertainty parameters (Root Mean Square (RMS), Variance, Skewness, Kurtosis, Hurst Exponent (Hurst, 1951), Shannon Entropy, Central Frequency, Maximum Frequency) are calculated for a given subject. Some of these parameters represent linear characteristics of the EEG signal and other represent non-linear properties of EEG (Diez, Torres, Avila, Laciar and Mut, 2009; Gupta and Agrawal, 2012; Gupta et al., 2015). In this work, the parameters are selected empirically as every signal or data has the distinguishable property in terms of a certain set of statistical parameters associated with the signal or data as shown in Figure 3.
4. Hence, final feature vector obtained after concatenation of features from six channels contains 192 parameters (4 IMFs corresponding to each segments × 8 parameters corresponding to each IMFs × 6 channels) for each task labels for a given subject.

**Figure 3:**
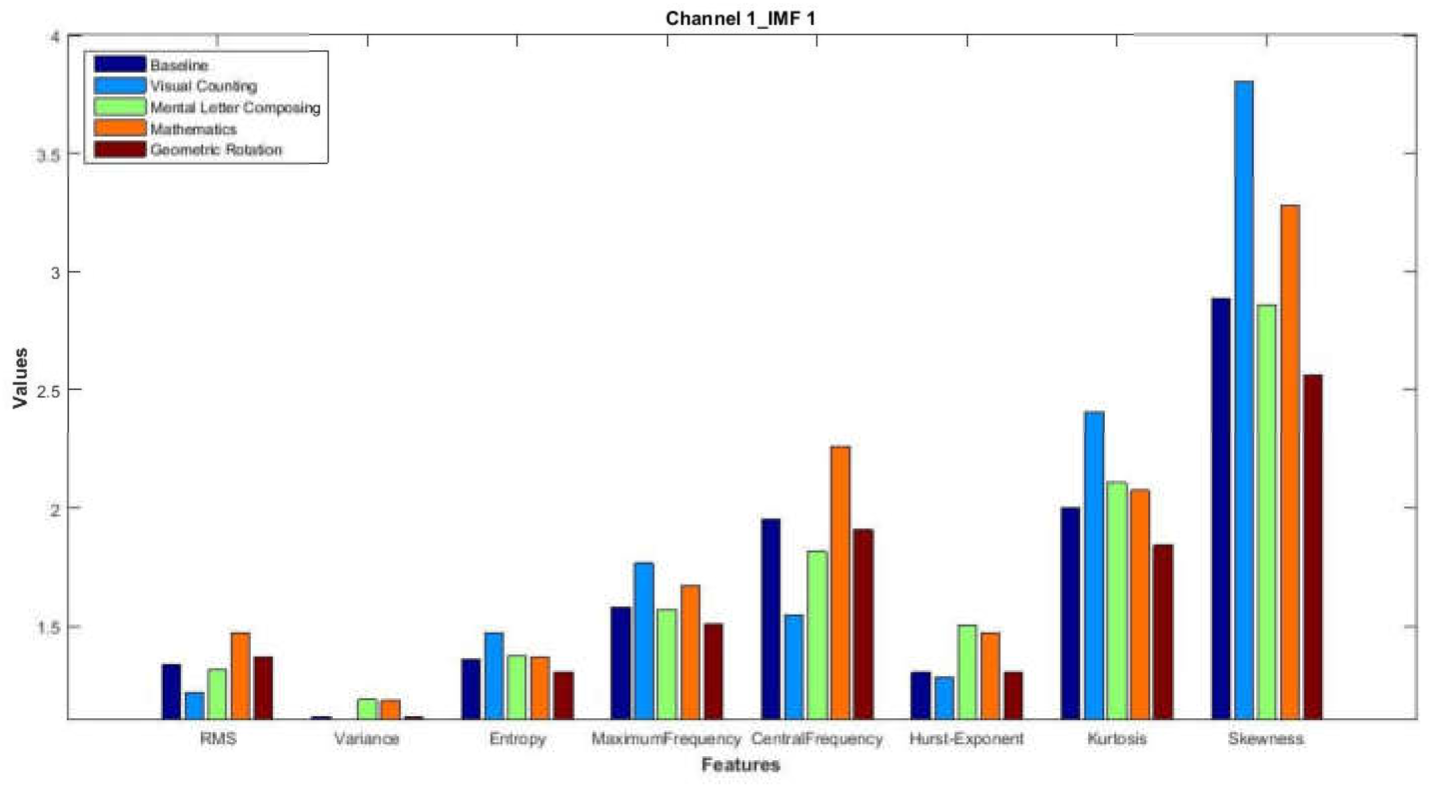
Eight features obtained corresponding to all five mental tasks for channel 1 from IMF 1 using EEMD method for Subject 1.

### 4.2 Result

The performance of the EMD and its variant has been evaluated in terms of classification accuracy achieved with SVM classifier with one versus all approach. Grid search is used to find optimal choice of regularization parameters. The average classification accuracy of 10 runs of 10 cross-validations is quoted. To check the efficacy of the proposed method, we have formulated three type of multi-mental task classification problems viz. three class, four class and five class as well as binary mental task classification.

#### Binary Class Problem

We have used binary combination of these tasks as BC, BL, BM, BR, CL, CM, CR, LM, LR and MR in this work.

#### Three Class Problem

In this problem, we have formed three-class mental tasks problems by choosing three different mental tasks at a time from given five mental tasks. There are ten different triplet mental task combinations for forming three class problem given as: BCL, BCM, BCR, BLM, BLR, BMR,CLM, CLR, CMR and LMR.

#### Four Class Problem

Construction of four mental task classification problem has been done by choosing four tasks at a time from the given five tasks. There are five different four class problems namely BCLM, BCLR, BCMR, BLMR and CLMR.

#### Five Class Problem

For the formation of the five mental task classification problem, we have taken all five mental tasks at a time. Thus, we have the five-class mental tasks classification problem as BCLMR.

Table 1 to Table 3 show the classification accuracy for the binary mental tasks classification problem of three different EMDs algorithms. The bold values show the best and average classification accuracy for different subjects. From these Tables, it is clear that among three EMDs algorithms, EEMD performs best for binary MTC. Similar kind of observation can be seen for three class, four class and five class of MTC, which have been shown from Table 4 to Table 10 respectively.

**Table 1:** Classification accuracy of EMD for binary mental task classification.

| Task-Combination | Sub 1 | Sub 2 | Sub 3 | Sub 5 | Sub 6 | Sub 7 | Average |
| --- | --- | --- | --- | --- | --- | --- | --- |
| BC | 92.33 | 77.74 | 72.35 | 63.18 | 86.80 | 84.47 | 79.48 |
| BL | 84.35 | 65.85 | 77.50 | 61.47 | 67.33 | 77.00 | 72.25 |
| BM | 92.93 | 87.40 | 76.45 | 70.85 | 89.25 | 92.10 | 84.83 |
| BR | 96.78 | **98.35** | 66.05 | 75.92 | 88.60 | 99.05 | 87.46 |
| CL | 68.45 | 77.79 | **84.15** | 67.02 | 78.03 | 92.16 | 77.93 |
| CM | 96.50 | 83.05 | 66.85 | 77.50 | **98.78** | 93.26 | 85.99 |
| CR | 74.65 | 90.21 | 58.35 | **80.38** | 87.18 | 99.32 | 81.68 |
| LM | 98.25 | 92.15 | 81.58 | 74.32 | 87.25 | 98.95 | **88.75** |
| LR | 86.98 | 97.65 | 75.60 | 75.27 | 81.13 | **99.45** | 86.01 |
| MR | **97.75** | 88.35 | 67.50 | 79.40 | 84.55 | 82.25 | 83.30 |
| Average | **88.90** | **85.85** | **72.64** | **72.53** | **84.89** | **91.80** | **82.77** |

**Table 2:** Classification accuracy of EEMD for binary mental task classification.

| Task-Combination | Sub 1 | Sub 2 | Sub 3 | Sub 5 | Sub 6 | Sub 7 | Average |
| --- | --- | --- | --- | --- | --- | --- | --- |
| BC | 93.75 | 90.85 | 72.33 | 94.42 | 89.85 | 91.65 | 88.81 |
| BL | 86.48 | 71.55 | 79.53 | 82.65 | 71.03 | 82.70 | 78.99 |
| BM | 93.23 | 88.95 | 80.33 | **97.15** | 94.23 | 96.80 | 91.78 |
| BR | 96.83 | 98.20 | 68.70 | 96.70 | 93.98 | 98.80 | **92.20** |
| CL | 71.30 | 88.50 | **85.03** | 69.92 | 82.45 | 91.90 | 81.52 |
| CM | 96.63 | 86.90 | 65.63 | 76.35 | **99.43** | 96.40 | 86.89 |
| CR | 76.60 | 95.35 | 60.60 | 81.35 | 92.00 | 98.55 | 84.08 |
| LM | **98.25** | 94.30 | 82.88 | 73.52 | 91.75 | 98.30 | 89.83 |
| LR | 87.00 | **98.95** | 77.50 | 76.13 | 89.03 | **100.00** | 88.10 |
| MR | 97.73 | 90.15 | 62.58 | 80.37 | 87.73 | 87.50 | 84.34 |
| Average | **89.78** | **90.37** | **73.51** | **82.86** | **89.15** | **94.26** | **86.65** |

**Table 3:** Classification accuracy of CEEMDAN for binary mental task classification.

| Task-Combination | Sub 1 | Sub 2 | Sub 3 | Sub 5 | Sub 6 | Sub 7 | Average |
| --- | --- | --- | --- | --- | --- | --- | --- |
| BC | 93.13 | 90.05 | 72.73 | 66.08 | 90.63 | 88.30 | 83.48 |
| BL | 86.20 | 71.30 | 78.93 | 62.85 | 73.10 | 81.00 | 75.56 |
| BM | 92.25 | 90.50 | 80.63 | 73.83 | 94.35 | 91.90 | 87.24 |
| BR | 97.60 | 99.20 | 67.73 | 78.40 | 94.08 | 98.25 | 89.21 |
| CL | 72.53 | 83.80 | **85.23** | 71.03 | 85.03 | 91.40 | 81.50 |
| CM | 97.03 | 87.20 | 67.63 | 75.47 | **99.68** | 95.30 | 87.05 |
| CR | 78.10 | 95.15 | 61.70 | 81.20 | 90.58 | 98.50 | 84.20 |
| LM | 97.43 | 93.45 | 81.38 | 73.70 | 92.10 | 98.75 | **89.47** |
| LR | 87.48 | **99.70** | 73.83 | 76.13 | 89.95 | **99.50** | 87.76 |
| MR | **98.18** | 90.60 | 64.50 | **81.38** | 88.80 | 84.30 | 84.63 |
| Average | **89.99** | **90.10** | **73.43** | **74.01** | **89.83** | **92.72** | **85.01** |

**Table 4:** Classification accuracy of EMD for three class mental task classification.

| Task-Combination | Sub 1 | Sub 2 | Sub 3 | Sub 5 | Sub 6 | Sub 7 | Average |
| --- | --- | --- | --- | --- | --- | --- | --- |
| BCL | 61.67 | 59.34 | 66.72 | 51.50 | 64.35 | 70.24 | 62.30 |
| BCM | 87.38 | 72.41 | 56.80 | 56.83 | **82.02** | 83.41 | 73.14 |
| BCR | 76.82 | 74.21 | 51.05 | 57.93 | 76.30 | 83.48 | 69.96 |
| BLM | 81.22 | 66.87 | **66.87** | 54.50 | 66.72 | 78.63 | 69.13 |
| BLR | 74.67 | 71.17 | 61.98 | 58.62 | 66.28 | 82.53 | 69.21 |
| BMR | **92.53** | 82.90 | 56.07 | 64.66 | 76.32 | 80.97 | **75.57** |
| CLM | 75.00 | 74.03 | 62.60 | 61.64 | 75.00 | 86.00 | 72.38 |
| CLR | 62.25 | 73.83 | 56.62 | 63.90 | 71.02 | **86.83** | 69.07 |
| CMR | 80.07 | 79.45 | 49.07 | **66.76** | 78.67 | 80.93 | 72.49 |
| LMR | 87.92 | **84.07** | 60.15 | 63.83 | 72.12 | 83.83 | 75.32 |
| Average | **77.95** | **73.83** | **58.79** | **60.02** | **72.88** | **81.69** | **70.86** |

**Table 5:** Classification accuracy of EEMD for three class mental task classification.

| Task-Combination | Sub 1 | Sub 2 | Sub 3 | Sub 5 | Sub 6 | Sub 7 | Average |
| --- | --- | --- | --- | --- | --- | --- | --- |
| BCL | 65.15 | 68.57 | 69.17 | 76.24 | 69.20 | 78.50 | 71.14 |
| BCM | 87.75 | 82.30 | 57.98 | 79.78 | **87.88** | 88.00 | 80.62 |
| BCR | 80.70 | 83.63 | 53.93 | 82.77 | 83.62 | 90.17 | 79.14 |
| BLM | 84.05 | 68.27 | **71.07** | 77.84 | 75.72 | 83.47 | 76.74 |
| BLR | 77.85 | 76.07 | 64.98 | 80.04 | 73.28 | 85.47 | 76.28 |
| BMR | **93.00** | 83.17 | 56.18 | **83.92** | 85.15 | 84.10 | **80.92** |
| CLM | 77.78 | 81.27 | 62.50 | 62.77 | 81.62 | **92.00** | 76.32 |
| CLR | 66.65 | 81.53 | 59.57 | 65.49 | 80.47 | 90.80 | 74.08 |
| CMR | 82.65 | 81.87 | 46.88 | 66.51 | 86.02 | 86.03 | 74.99 |
| LMR | 88.32 | **88.37** | 58.18 | 64.90 | 81.60 | 86.00 | 77.89 |
| Average | **80.39** | **79.50** | **60.05** | **74.03** | **80.46** | **86.45** | **76.81** |

**Table 6:** Classification accuracy of CEEMDAN for three class mental task classification.

| Task-Combination | Sub 1 | Sub 2 | Sub 3 | Sub 5 | Sub 6 | Sub 7 | Average |
| --- | --- | --- | --- | --- | --- | --- | --- |
| BCL | 64.58 | 67.77 | 69.38 | 51.84 | 69.30 | 77.50 | 66.73 |
| BCM | 86.63 | 82.87 | 58.27 | 56.71 | **87.22** | 84.93 | 76.10 |
| BCR | 80.90 | 81.20 | 52.65 | 60.24 | 84.35 | 88.53 | 74.65 |
| BLM | 83.42 | 68.67 | **69.95** | 54.79 | 76.30 | 82.93 | 72.68 |
| BLR | 77.63 | 75.00 | 64.45 | 57.57 | 72.83 | 87.43 | 72.49 |
| BMR | **92.60** | 85.97 | 57.18 | 67.00 | 84.53 | 80.33 | **77.94** |
| CLM | 77.62 | 78.73 | 62.13 | 62.84 | 82.30 | 89.37 | 75.50 |
| CLR | 66.42 | 76.30 | 58.85 | 66.56 | 79.57 | **90.63** | 73.05 |
| CMR | 83.32 | 84.90 | 49.27 | **67.38** | 85.75 | 85.43 | 76.01 |
| LMR | 88.32 | **87.70** | 57.52 | 64.70 | 82.88 | 85.23 | 77.73 |
| Average | **80.14** | **78.91** | **59.97** | **60.96** | **80.50** | **85.23** | **74.29** |

**Table 7:** Classification accuracy of EMD for four class mental task classification.

| Task-Combination | Sub 1 | Sub 2 | Sub 3 | Sub 5 | Sub 6 | Sub 7 | Average |
| --- | --- | --- | --- | --- | --- | --- | --- |
| BCLM | 65.03 | 57.28 | **55.18** | 49.81 | 63.00 | 70.92 | 60.20 |
| BCLR | 66.80 | **68.21** | 48.19 | **56.21** | 64.46 | **76.97** | 63.47 |
| BCMR | 74.96 | 65.08 | 53.36 | 53.33 | 60.95 | 72.00 | 63.28 |
| BLMR | **76.48** | 67.72 | 45.76 | 54.80 | **71.01** | 74.64 | **65.07** |
| CLMR | 56.50 | 58.59 | 48.64 | 52.48 | 60.90 | 71.13 | 58.04 |
| Average | **67.95** | **63.37** | **50.23** | **53.33** | **64.07** | **73.13** | **62.01** |

**Table 8:** Classification accuracy of EEMD for four class mental task classification.

| Task-Combination | Sub 1 | Sub 2 | Sub 3 | Sub 5 | Sub 6 | Sub 7 | Average |
| --- | --- | --- | --- | --- | --- | --- | --- |
| BCLM | 69.54 | 65.05 | **57.11** | 67.96 | 71.08 | 78.55 | 68.21 |
| BCLR | 71.40 | **75.95** | 46.85 | 57.10 | 75.93 | **83.63** | 68.48 |
| BCMR | 77.36 | 68.73 | 54.25 | 69.86 | 71.28 | 78.20 | 69.95 |
| BLMR | **78.60** | 75.80 | 45.16 | **70.50** | **79.00** | 79.70 | **71.46** |
| CLMR | 61.66 | 65.60 | 52.76 | 69.70 | 68.40 | 80.53 | 66.44 |
| Average | **71.71** | **70.23** | **51.23** | **67.02** | **73.14** | **80.12** | **68.91** |

**Table 9:** Classification accuracy of CEEMDAN for four class mental task classification.

| Task-Combination | Sub 1 | Sub 2 | Sub 3 | Sub 5 | Sub 6 | Sub 7 | Average |
| --- | --- | --- | --- | --- | --- | --- | --- |
| BCLM | 67.63 | 63.73 | **55.36** | 49.09 | 70.66 | 74.90 | 63.56 |
| BCLR | 69.23 | 74.13 | 47.91 | **57.83** | 75.98 | **80.98** | 67.67 |
| BCMR | 77.48 | 69.70 | 54.48 | 52.44 | 72.05 | 76.53 | 67.11 |
| BLMR | **78.05** | **76.90** | 45.64 | 55.27 | **79.16** | 76.65 | **68.61** |
| CLMR | 60.55 | 64.25 | 53.00 | 50.52 | 67.28 | 79.40 | 62.50 |
| Average | **70.59** | **69.74** | **51.28** | **53.03** | **73.03** | **77.69** | **65.89** |

**Table 10:** Classification accuracy for all five class mental task classification of all feature extraction method.

| Task-Combination | Feature Extraction methods | Sub 1 | Sub 2 | Sub 3 | Sub 5 | Sub 6 | Sub 7 | Average |
| --- | --- | --- | --- | --- | --- | --- | --- | --- |
| BCLMR | EMD | 59.60 | 56.71 | 44.53 | 53.26 | 57.47 | 66.41 | 56.33 |
|  | EEMD | **65.23** | **63.00** | 44.69 | **62.04** | 67.47 | **74.26** | **62.78** |
|  | CEEMDAN | 63.85 | 62.92 | **46.93** | 48.45 | **67.81** | 71.40 | 60.23 |

### 4.3 Comparison of the proposed model for multi mental task classification problem

In this subsection, we have discussed and compared the proposed approach with the work of Zhang et al. (2010) multi mental task classification. Table 11 shows the comparison of the work of Zhang et al. (2010) for multi-mental task classification.

**Table 11:** Comparison table of classification accuracy achieved for multi mental task classification of the work of Zhang et al. (2010) with proposed approach.

|  | Two class classification |  |  | Three class classification |  |  | Four class classification |  |  | Five class classification |  |  |
| --- | --- | --- | --- | --- | --- | --- | --- | --- | --- | --- | --- | --- |
| Zhang et al. (2010) | A | B | C | A | B | C | A | B | C | A | B | C |
| Sub1 | 77.60 | **85.90** | 83.80 | 63.90 | **75.30** | 70.90 | 54.40 | **66.60** | 60.50 | 47.60 | **60.40** | 55.40 |
| Sub2 | 62.90 | **67.50** | 66.20 | 46.50 | **53.80** | 47.90 | 37.90 | **45.40** | 38.30 | 31.90 | **39.90** | 33.60 |
| Sub3 | 69.40 | **72.50** | 71.50 | 54.10 | **59.40** | 57.00 | 45.30 | **52.10** | 49.80 | 39.30 | **46.30** | 43.70 |
| Proposed approach | EMD | EEMD | CEEMDAN | EMD | EEMD | CEEMDAN | EMD | EEMD | CEEMDAN | EMD | EEMD | CEEMDAN |
| Sub1 | 88.90 | 89.78 | **89.99** | 77.95 | **80.39** | 80.14 | 67.95 | **71.71** | 70.59 | 59.60 | **65.23** | 63.85 |
| Sub2 | 85.85 | **90.37** | 90.10 | 73.83 | **79.50** | 78.91 | 63.37 | **70.23** | 69.74 | 56.71 | **63.00** | 62.92 |
| Sub3 | 72.64 | **73.51** | 73.43 | 58.79 | **60.05** | 59.97 | 50.23 | 51.23 | **51.28** | 44.53 | 44.69 | **46.93** |

In the Table 11, methods A, B and C are the schemes used by Zhang et al. (2010) based on asymmetry ratio for calculation of different number of frequency band powers using 75-dimensional, 90-dimensional and 42-dimensional feature vector, respectively. From this Table, it is clear that our approach for creating features vectors outperforms in terms of average classification accuracy for all the three subjects for all the multi mental tasks classification problem.

### 4.4 Discussion

Since EEG signal have non-linear and non-stationary property, thus there is a need of an algorithm which can capture such properties of the signal. EMD is such an algorithm which can capture tempo-spectral information of the signal. After decomposing the signal into high and low frequency components, it is important to extract some statistical and uncertainty parameters from this decomposed signal for compact representation in terms of features which can help in differentiating one mental state to another. In addition, there are two improved version of EMD algorithm named as EEMD and CEEMDAN algorithm, which can capture tempo-spectral information even from noise assisted signal.

Figure 4 to Figure 7 represent the average classification over all tasks combination for all the possible combination of mental tasks of all subjects. From these Figures, it is clear that EEMD algorithm outperforms when compared to EMD and CEEMDAN. It is also observed that for the Sub 1, Sub 2 and Sub 7, the distinguishing capacity of the classification model to differentiate the two or more mental tasks simultaneously is better than oth r subjects, from the extracted featur s by the EMDs algorithms.

**Figure 4:**
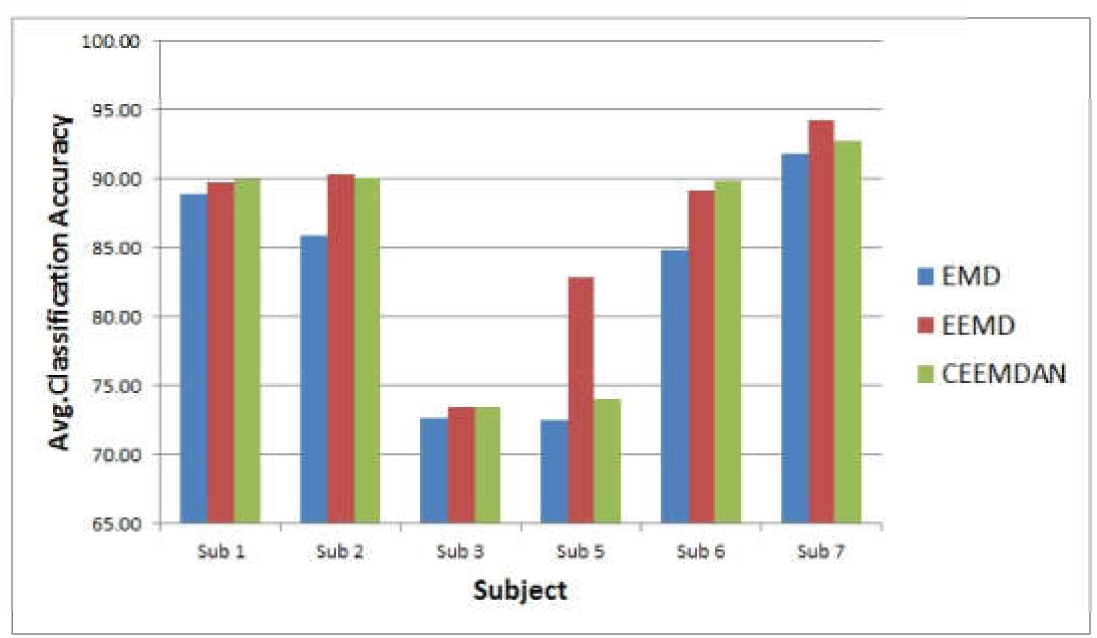
Bar chart for the average classification accuracy over all binary mental tasks for all six subjects.

**Figure 5:**
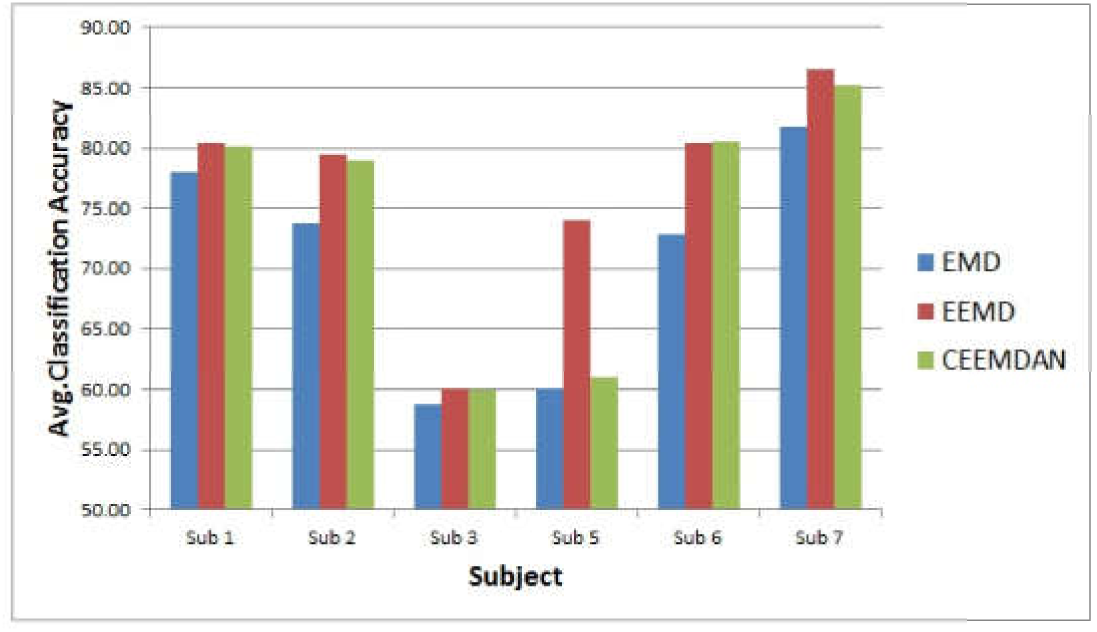
Bar chart for the average classification accuracy over all three class mental tasks for all six subjects.

**Figure 6:**
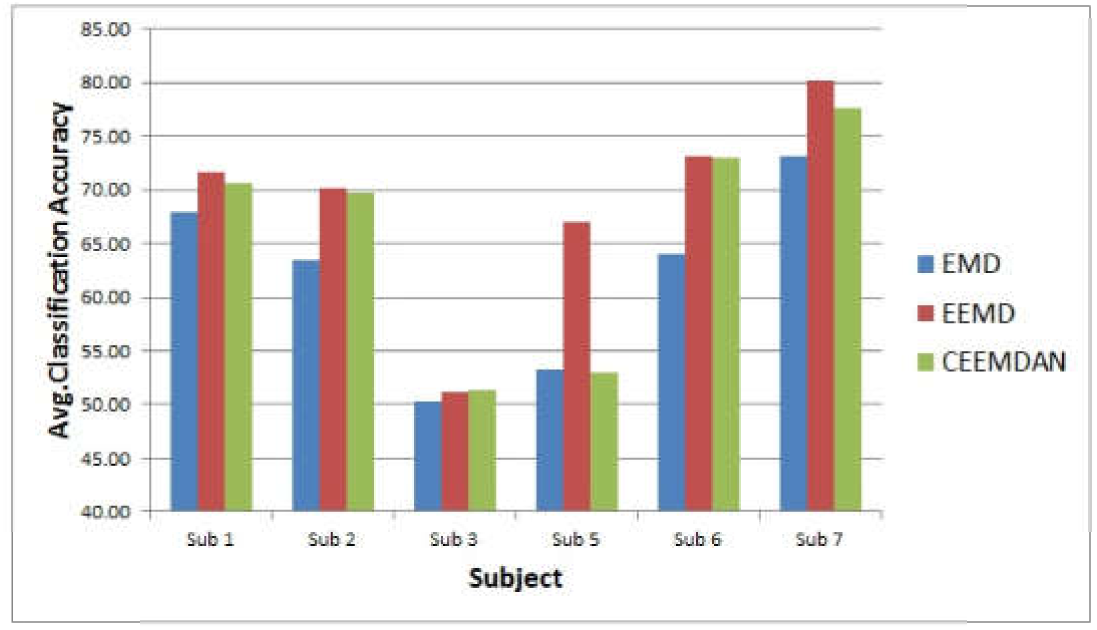
Bar chart for the average clas ification accuracy over all four class mental tasks for all six subjects.

**Figure 7:**
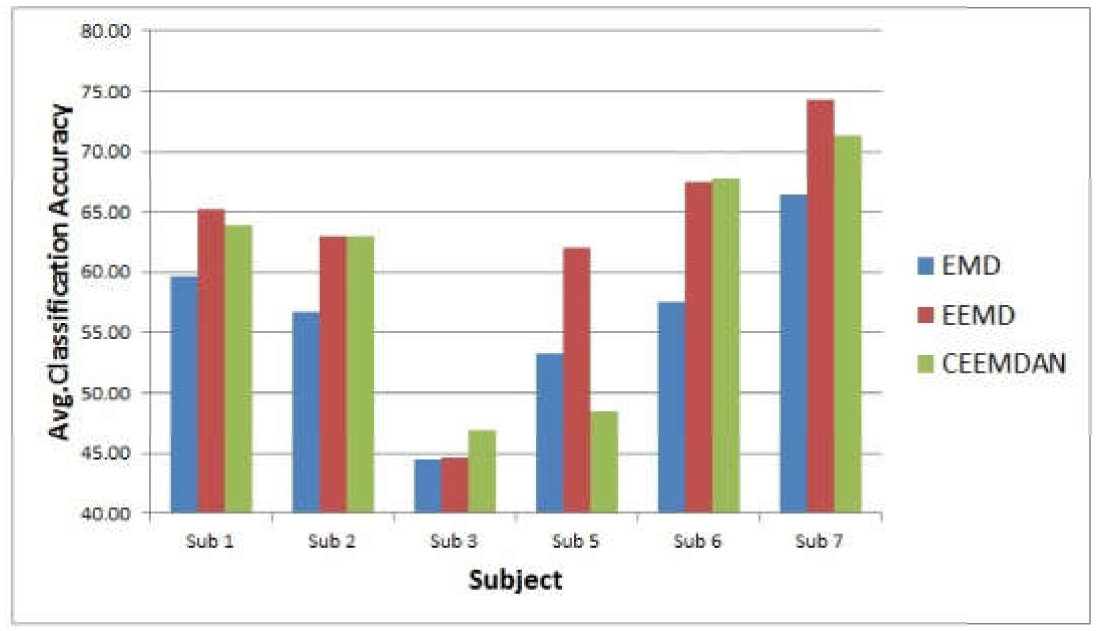
Bar chart for the average classification accuracy over all five class mental tasks for all six subjects.

### 4.5 Statistical Test

We have utilized a two way, non-parametric statistical test known as Friedman test (Derrac et al., 2011; Friedman, 1937) to find out the significant difference among these three EMD methods for EEG signal. The Table 12 shows the average Friedman ranking of the methods for different combination of metal tasks classification problem, which shows that EEMD method outperform among three methods for all the possible metal tasks classification problem.

The performance of different EMD methods (in this work) is studied with respect to control method i.e. best performer from the Friedman’s ranking (which is EEMD). The test statistics for the comparison of *m*^*th*^ method to *n*^*th*^ method, *z*, is given as

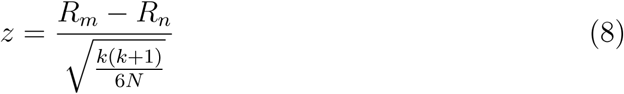

where *R*_*m*_ and *R*_*n*_ are the average ranking of the methods, *k* and *N* are the number of methods (algorithms) and experiments respectively. However, these *p* values so obtained are not suitable for comparison with the control method. Instead, adjusted p values (Derrac et al., 2011) are computed which take into account the error accumulated and provide the correct correlation. For this, a set of post-hoc procedures are defined and adjusted *p* values are computed. For pair-wise comparisons, the widely used post-hoc methods to obtain adjusted *p* values are (Derrac et al., 2011): Bonferroni-Dunn, Holm, Hochberg and Hommel procedures. Table 13 shows the various value of adjusted *p* values obtained from aforementioned methods. From this Table, it is clear that there is statistical difference between EEMD and other two methods.

**Table 12:** Average Rankings of the algorithms

| Algorithm | Ranking |  |  |  |
| --- | --- | --- | --- | --- |
| method | Binary Class | Three Class | Four Class | Five Class |
| EMD | 3.00 | 3.00 | 3.00 | 2.93 |
| **EEMD** | **1.03** | **1.01** | **1.03** | **1.17** |
| CEEMDAN | 1.97 | 1.99 | 1.97 | 1.90 |

## 5 Conclusion

Classification of EEG signal for any purpose requires detailed analysis of the signal, i.e. intrinsic properties of the signal. This work presented a comprehensive study of the variants of EMD algorithm to find intrinsic characteristics of the EEG signal for mental task classification. After decomposing the signal through the EMDs algorithms, 8 parameters were calculated from each segment of the decomposed signal to form the feature vector from the signal. SVM is utilized for classification purpose. Experimental results showed that EEMD algorithm perform best among three EMD algorithms. Statistical analysis are also performed to investigate whether three EMD algorithms statistically different or not for MTC.

**Table 13:** Adjusted *p*-values

| Class Combinations | Algorithm | unadjusted $p$ | $p_{Bonf}$ | $p_{Holm}$ | $p_{Hoch}$ | $p_{Hommel}$ |
| --- | --- | --- | --- | --- | --- | --- |
| Binary Class | EMD | 4.16E-44 | 8.33E-44 | 8.33E-44 | 8.33E-44 | 8.33E-44 |
|  | CEEMDAN | 2.99E-11 | 5.99E-11 | 2.99E-11 | 2.99E-11 | 2.99E-11 |
| Three Class | EMD | 5.69E-45 | 1.14E-44 | 1.14E-44 | 1.14E-44 | 1.14E-44 |
|  | CEEMDAN | 4.22E-12 | 8.44E-12 | 4.22E-12 | 4.22E-12 | 4.22E-12 |
| Four Class | EMD | 4.16E-44 | 8.33E-44 | 8.33E-44 | 8.33E-44 | 8.33E-44 |
|  | CEEMDAN | 2.99E-11 | 5.99E-11 | 2.99E-11 | 2.99E-11 | 2.99E-11 |
| Five Class | EMD | 1.49E-35 | 2.97E-35 | 2.97E-35 | 2.97E-35 | 2.97E-35 |
|  | CEEMDAN | 2.44E-7 | 4.89E-7 | 2.44E-7 | 2.44E-7 | 2.44E-7 |

In the future work, we would like to explore some advanced decomposition methods for the EEG signal. To further reduce the dimensionality Feature selection approach can be investigated to improve the classification performance for MTC. It is also interesting to investigate some new set of parameters associated with the signals which can help in distinguishing different mental states more accurately.

## Acknowledgment

The authors of this manuscript express their gratitude to the scheme Cognitive Science Research Initiative (CSRI), Department of Science & Technology, Ministry of Science & Technology, Govt. of India. The authors are also thankful to the grant number BT/BI/03/004/2003(C) Bioinformatics division,Department of Biotechnology, Ministry of Science & Technology, Govt. of India.

## Footnotes

1 http://www.cs.colostate.edu/eeg/main/data/1989_Keirn_and_Aunon

